# 7SK methylation Promotes Transcriptional Activity

**DOI:** 10.1101/2022.10.17.512631

**Authors:** Marcelo Perez-Pepe, Anthony W. Desotell, Hengyi Li, Wenxue Li, Bing Han, Qishan Lin, Daryl E. Klein, Yansheng Liu, Hani Goodarzi, Claudio R. Alarcón

**Affiliations:** Department of Pharmacology, Yale University School of Medicine, New Haven, CT 06520, USA; Yale Cancer Biology Institute, Yale University, West Haven, CT 06516, USA; RNA Epitranscriptomics and Proteomics Resource, University at Albany, Albany, NY 12222, USA; Department of Biochemistry and Biophysics, University of California, San Francisco, San Francisco, CA 94158, USA

## Abstract

A fundamental facet of cell signaling is the conversion of extracellular signals into adaptive transcriptional responses. The role of RNA modifications in this process is poorly understood. The small nuclear RNA 7SK prevents transcription elongation by sequestering the complex CDK9/CCNT1 (P-TEFb). We discovered that METTL3 methylates 7SK. The m^6^A methylation of 7SK in turn promotes its binding to heterogeneous nuclear ribonucleoproteins (HNRNPs), with consequent release of the HEXIM1/P-TEFb complex – leading to the induction of growth factor-stimulated transcriptional responses. The methylation of 7SK relies on the activation of METTL3 via phosphorylation downstream of growth factors-signaling pathways such as the epidermal growth factor (EGF). Our findings establish a novel function for the m^6^A modification in converting growth-factor signaling events to a transcriptional elongation regulatory response via an RNA-methylation-dependent switch.

**One-Sentence Summary:** m^6^A methylation of the non-coding RNA 7SK promotes transcriptional activity upon growth factor stimulation.

---

Extracellular signals regulate intracellular pathways that control adaptive responses such as growth, quiescence, differentiation, and cell migration. Ultimately, these signaling pathways converge on the nucleus to elicit specific transcriptional programs. 7SK (RN7SK) is a 332-nucleotide (nt) small nuclear RNA (snRNA) that interacts with multiple RNA-binding proteins to regulate transcriptional elongation. 7SK sequesters the transcription elongation factors CDK9/CCNT1 (P-TEFb) by interacting with the protein HEXIM1 (*1, 2*). Under conditions favoring cell proliferation, 7SK dissociates from HEXIM1/P-TEFb and binds to HNRNP proteins, including HNRNPA1, HNRNPA2B1, HNRNPR, and HNRNPQ (SYNCRIP) (*3, 4*). Once P-TEFb is released from 7SK, it phosphorylates serine 2 of the RNA Pol II C-terminal domain (CTD), allowing transcription elongation to proceed (*5, 6*).

RNA modifications are a key feature of post-transcriptional gene expression control. Methylation of RNA on the *N*^6^ nitrogen of adenosine (m^6^A) is the most abundant post-transcriptional modification in messenger RNA (mRNA) (*7*). We and others have implicated m^6^A, and the enzyme responsible for RNA methylation METTL3 (*8-10*), in regulation of RNA stability (*11*), miRNA processing (*12, 13*), RNA splicing (*12, 14*), and translation (*15*). These processes impact cellular functions such as meiosis (*16, 17*), cell proliferation (*18, 19*) and embryonic stem cell differentiation (*19*), and pathophysiological states such as cancer (*20, 21*). Despite the known general roles for m^6^A in cellular homeostasis (*21-24*), it remains unclear how signaling pathways impact METTL3 function and downstream transcriptional activity.

## Results

### METTL3 methylates the snRNA 7SK

Nuclear m^6^A-seq experiments revealed the presence of m^6^A modifications on 7SK (Fig. 1A). Furthermore, high-throughput sequencing of RNA isolated in crosslinking immunoprecipitation (HITS-CLIP) of METTL3, revealed direct binding of METTL3 to the 7SK RNA (Fig. 1A) (*12*) – suggesting that METTL3 may recognize and methylate 7SK. To further validate these results with an independent method, we pulled down endogenous 7SK and used RNA mass spectrometry to assess m^6^A content. To isolate endogenous 7SK, we took advantage of its known interaction with the protein LARP7 (*25, 26*). LARP7 binds 7SK constitutively thus protecting it from degradation (*27-30*), which contrasts with the binding of HEXIM1/P-TEFb or HNRNPs to 7SK, which depends on the cell state. We therefore generated a HeLa cell line stably expressing low levels of Flag-tagged LARP7 protein. To facilitate the isolation of native LARP7-7SK complexes, we performed mechanical disruption of cells at cryogenic temperature (cryomilling) using liquid nitrogen, thus preserving post-translational states and high order complexes (*31*). As expected, immunoprecipitation of Flag-tagged LARP7 (Flag-IP) efficiently pulled down endogenous 7SK as the main RNA species, as revealed by acrylamide gel electrophoresis and Northern blotting (Fig. 1B). We extracted 7SK from the gel and, after purification, digested it with a mixture of enzymes to generate single nucleotides. Levels of m^6^A and adenosine were then measured by ultra-performance liquid chromatography coupled with tandem mass spectrometry (UHPLC-MS/MS). Mass spectrometry detected the presence of m^6^A in endogenous 7SK, thus validating its methylation status and excluding the possibility of cross-recognition of the anti-m^6^A antibody with other potential RNA modifications (Fig. 1C and S1B).

**Fig. 1.**
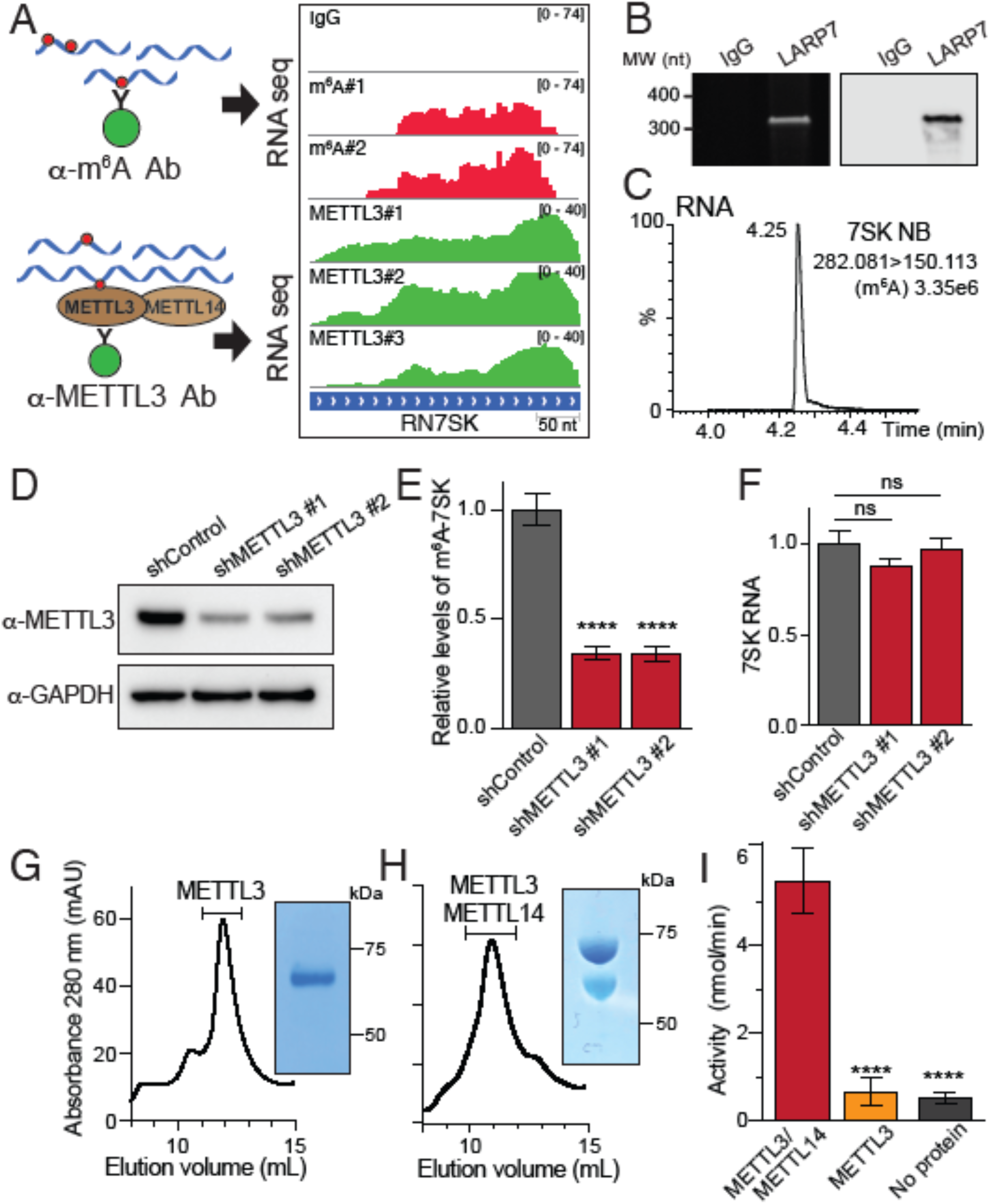
7SK is m^6^A methylated by METTL3. **(A)** Tracks of m^6^A-seq and METTL3 HITS-CLIP on 7SK (right). m^6^A-seq experiments were done in biological duplicates and METTL3 HITS-CLIP in biological triplicates. The top panel shows an IgG negative control. On the left, a schematic representation of m^6^A-seq (top) and METTL3 HITS-CLIP (bottom) experiments. **(B)** Isolation of endogenous 7SK by immunoprecipitation of Flag-tagged LARP7 using an anti-Flag antibody. Purified endogenous 7SK as shown by a urea-acrylamide gel (left) and Northern blot analysis (right). Samples using IgG for IP are shown as controls. **(C)** Detection of m^6^A in endogenous 7SK through UHPLC-MS/MS from the RNA isolated in 1B. The figure shows a representative spectrum from two biological replicates, with three technical replicates each. **(D)** Knockdown of METTL3 using two independent shRNAs. Total cell extract was analyzed by Western blot using the indicated antibodies. Blots are a representation of three biological repeats. **(E)** m^6^A methylated nuclear RNA was immunoprecipitated and the levels of 7SK were quantified by qRT-PCR. The bar graph depicts the effect of METTL3 depletion using two independent shRNAs on the levels of methylated 7SK. Total endogenous 7SK levels were used for normalization. The graph shows mean ± SD from a representative experiment out of two biological replicates and three technical replicates each. One-way ANOVA, Dunnett’s post-test, \*\*\*\**P*<1×10^−4^. **(F)** Levels of 7SK were quantified by qRT-PCR. The bar graph depicts the effect of METTL3 depletion using two independent shRNAs. The graph shows mean ± SD from a representative experiment out of two biological replicates and three technical replicates each. One-way ANOVA, Dunnett’s post-test. **(G**,**H)** METTL3 (G) and METTL3/METTL14 protein complexes (H) were expressed in insect cells, affinity-purified using Ni-beads, and separated using size exclusion chromatography (SEC). Elution profiles of the proteins after separation through a Superdex 200 Increase 10/300 GL column. The inset shows a representative image of a Coomassie staining of SDS-PAGE from one of the fractions corresponding to the peak (indicated in the chromatogram). (**I)** Methylation activity of the METTL3/METTL14 complex and METTL3 alone using *in vitro* transcribed 7SK as a substrate. METTL3/METTL14 and METTL3 proteins were expressed in insect cells, affinity purified, and separated by SEC. A reaction without protein was used as a negative control for the assay. The graph shows mean ± SD from a representative experiment out of three biological replicates, with three technical replicates each. One-way ANOVA, Dunnett’s post-test, \*\*\*\**P*<1×10^−4^.

To confirm that this methylation is mediated by METTL3, we depleted METTL3 by RNAi and quantified the effect on abundance of methylated 7SK. To do this we generated stable cell lines expressing two independent shRNAs against METTL3 and performed an m^6^A immunoprecipitation of nuclear RNA followed by qRT-PCR of 7SK. Depleting METTL3 reduced methylated 7SK levels by almost 3-fold, without affecting total 7SK levels (Fig. 1, D-F). We next tested the ability of the purified METTL3/METTL14 complex (co-expressed in insect cells) to methylate *in vitro* transcribed (IVT) 7SK, which it was able to do efficiently (Fig. 1, G-I). Importantly, 7SK methylation required both METTL3 and METTL14, with METTL3 alone being insufficient to drive methylation (Fig. 1I). Together, these results demonstrate that METTL3 methylates 7SK.

### m^6^A favors binding of HNRNP proteins to 7SK

Although the structure of 7SK is not fully elucidated and could be altered by binding of different protein complexes, secondary structure predictions and previous SHAPE experiments identify several distinct stem-loops in 7SK (*32-34*). HEXIM1 binds a stem-loop close to the 5’ end of the RNA, and the crystal structure of a partial HEXIM1 protein associated with a dsRNA fragment corresponding to 7SK nucleotides 24-87 has been determined (*35*). HNRNPA1, HNRNPA2B1, HNRNPQ, and HNRNPR mainly bind to a hairpin corresponding to 7SK nucleotides 196-277 (*33, 36*). LARP7 and MePCE protect 7SK from degradation by binding to the junction of the 5’ and 3’ ends (*27-30*) (Fig. 2A).

**Fig. 2.**
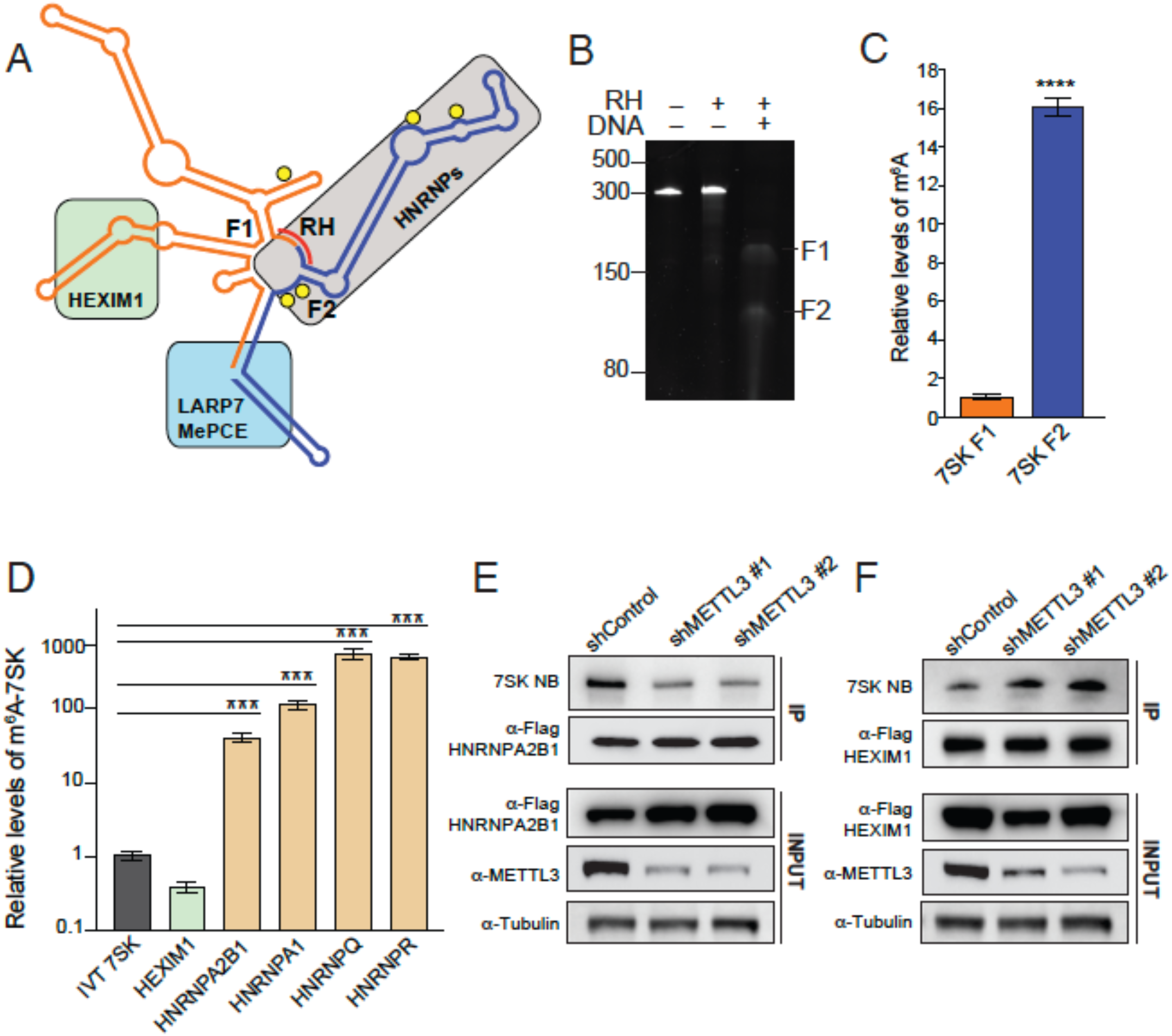
HEXIM1 preferentially binds unmethylated 7SK. **(A)** 7SK secondary structure representation. Hairpins known to be recognized by different proteins are depicted as a green box (HEXIM1/P-TEFb), grey (HNRNPs), and light blue (LARP7 and MePCE). Predicted m^6^A motifs are shown as yellow circles and the complementary DNA oligo used to fragment 7SK in the RNase H (RH) digestion is shown as a red line. **(B)** RNase H digestion of endogenous 7SK. TBE-urea acrylamide RNA gel showing untreated 7SK and treated with RNase H, with or without the addition of the complementary DNA oligo. **(C)** Relative m^6^A levels in the two fragments generated by RNase H treatment of endogenous 7SK pulled-down using Flag-tagged LARP7 as shown in B. After purification, 7SK was digested by RNase H using the DNA complementary oligo depicted in Fig. 1F. After fragmentation, the purified fragments were subjected to m^6^A-IP followed by qRT-PCR quantification. **(D)** Stably expressing Flag-tagged proteins were used to pull down endogenous 7SK. After purification of the pulled-down 7SK RNA, a second IP using an anti-m^6^A antibody was performed, and the levels of 7SK were quantified by qRT-PCR. Non-methylated *in vitro* transcribed 7SK RNA (IVT 7SK) was used as a negative control for the m^6^A IP. The graph shows mean ± SD from a representative experiment, out of two biological replicates, with three technical replicates each. One-way ANOVA, Dunnett’s post-test, \*\*\**P*<1×10^−3^. **(E-F)** 7SK-protein interaction with HNRNPA2B1 (E) and HEXIM1 (F) upon METTL3 depletion using two independent shRNAs. The top two panels represent the pull-down of endogenous 7SK measured by NB and the loading immunoprecipitated corresponding proteins detected by WB. The bottom three panels show the input protein levels before the IP. Blots are representative of at least two biological repeats.

Sequence analysis reveals that 7SK contains five canonical METTL3 recognition sequences (DRACH) located between nucleotides 170 and 290. Four of these motifs are located within the known hairpin that is recognized by HNRNP proteins (yellow circles in Fig. 2A). To better understand the location of m^6^A modifications, we fragmented 7SK and analyzed m^6^A levels in each fragment. We first pulled down endogenous 7SK using Flag-LARP7 and extracted 7SK from the complex. We then used a DNA oligo complementary to the sequence between positions 190 and 198. The hybridization of this DNA oligo will guide RNase H to cleave the resulting RNA-DNA hybrids and split 7SK into two fragments, F1 and F2 (Fig. 2, A and B). After RNase H cleavage, the fragments were isolated and subjected to m^6^A-IP. After the IP, the fragments were further purified and quantified by qRT-PCR. We found that the 3’ fragment (F2), which contains four of the five predicted m^6^A motifs, exhibited the highest levels of m^6^A (Fig. 2C).

To understand the effects of m^6^A modifications on 7SK, we tested the ability of the known 7SK-interacting proteins to interact with methylated 7SK. To do this we generated independent cell lines stably expressing the main 7SK direct-interactors (HEXIM1, CDK9, CCNT1, HNRNPA2B1, HNRNPA1, HNRNPQ, and HNRNPR) as Flag-tagged proteins in HeLa cells. The generation of these multiple cell lines facilitates the normalization of the immunoprecipitation protocol, making it less dependent on the intrinsic differences among various antibodies recognizing distinct epitopes in each ribonucleoprotein complex. After Flag-IPs of each independent protein, 7SK was extracted and purified and a second immunoprecipitation using an m^6^A antibody or control beads was performed. The methylated immunoprecipitated 7SK was then analyzed by qRT-PCR and normalized by the total amount of 7SK immunoprecipitated by the Flag-IP. As a negative control for m^6^A-IP, we used an IVT non-methylated 7SK. As shown in Fig. 2D, all the HNRNP proteins efficiently pulled down m^6^A-methylated 7SK between 50-500 fold more efficiently than the background IVT negative control. In contrast, HEXIM1 pulled down m^6^A-methylated 7SK at a level comparable to the IVT control (Fig. 2D), indicating that HNRNP proteins but not HEXIM1 interact with m^6^A-methylated 7SK. We then tested the impact of METTL3 depletion on the ability of these proteins to interact with endogenous 7SK. We depleted METTL3 from cells expressing Flag-HNRNPA2B1 as a representative of the HNRNP proteins and from cells expressing Flag-HEXIM1. As shown in Fig. 2E, METTL3 depletion reduced the levels of 7SK co-immunoprecipitated with HNRNPA2B1, but increased 7SK immunoprecipitation with HEXIM1 (Fig. 2F), while not affecting the association of 7SK with LARP7. These results demonstrate that the region of 7SK recognized by HNRNP proteins is the most m^6^A methylated. Moreover, this methylation facilitates the interaction of 7SK with HNRNPs and has the opposite effect on HEXIM1.

### EGF modulates the interaction between METTL3 and HEXIM1

The positive effect of m^6^A on cell proliferation is consistent with the formation of a complex between HNRNPs and m^6^A-7SK and the release of HEXIM1/P-TEFb to allow transcriptional elongation. Since pro-proliferative cell states favor formation of the 7SK-HNRNP complex and the release of HEXIM1/P-TEFb, we hypothesized that signaling pathways might elevate METTL3 activity. Post-translational modifications that stimulate methylation of 7SK would promote release of the transcription elongation factors to enhance proliferation. Since protein phosphorylation is a primary mechanism for regulating protein function in response to extra-and intracellular signals, we used immunoprecipitation coupled to phospho-mass spectrometry to investigate whether growth factor stimulation alters METTL3 phosphorylation. We first focused on the Epidermal Growth Factor (EGF) due to its established role in cell proliferation. EGF stimulation induced METTL3 phosphorylation at serine 43 (pS43). Based on these results, we generated polyclonal antibodies against pS43-METTL3 and validated the EGF-induced phosphorylation of METTL3 (Fig. 3A). Since pS43 is a predicted ERK1 phosphorylation site (Ser/Thr-Pro), we next tested the requirement of ERK for the S43 METTL3 phosphorylation. We used the general MEK inhibitor U0126 to block ERK activation. As shown in Fig. 3A, U0126 completely abolished EGF-induced ERK activation in HeLa cells and, importantly, prevented the EGF-dependent increase in S43-METTL3 phosphorylation (Fig. 3A). To determine whether ERK can phosphorylate METTL3 directly, we performed *in vitro* kinase assays using recombinant ERK1. We expressed and isolated human METTL3 from bacteria as a GST-fusion protein to eliminate endogenous phosphorylation events that may occur in eukaryotic cells. Incubation of ERK with METTL3 *in vitro* resulted in robust phosphorylation of METTL3 on S43 as measured by phospho-mass spectrometry. These results demonstrate that ERK phosphorylates METTL3 at S43 downstream of growth factor stimulation.

**Fig. 3.**
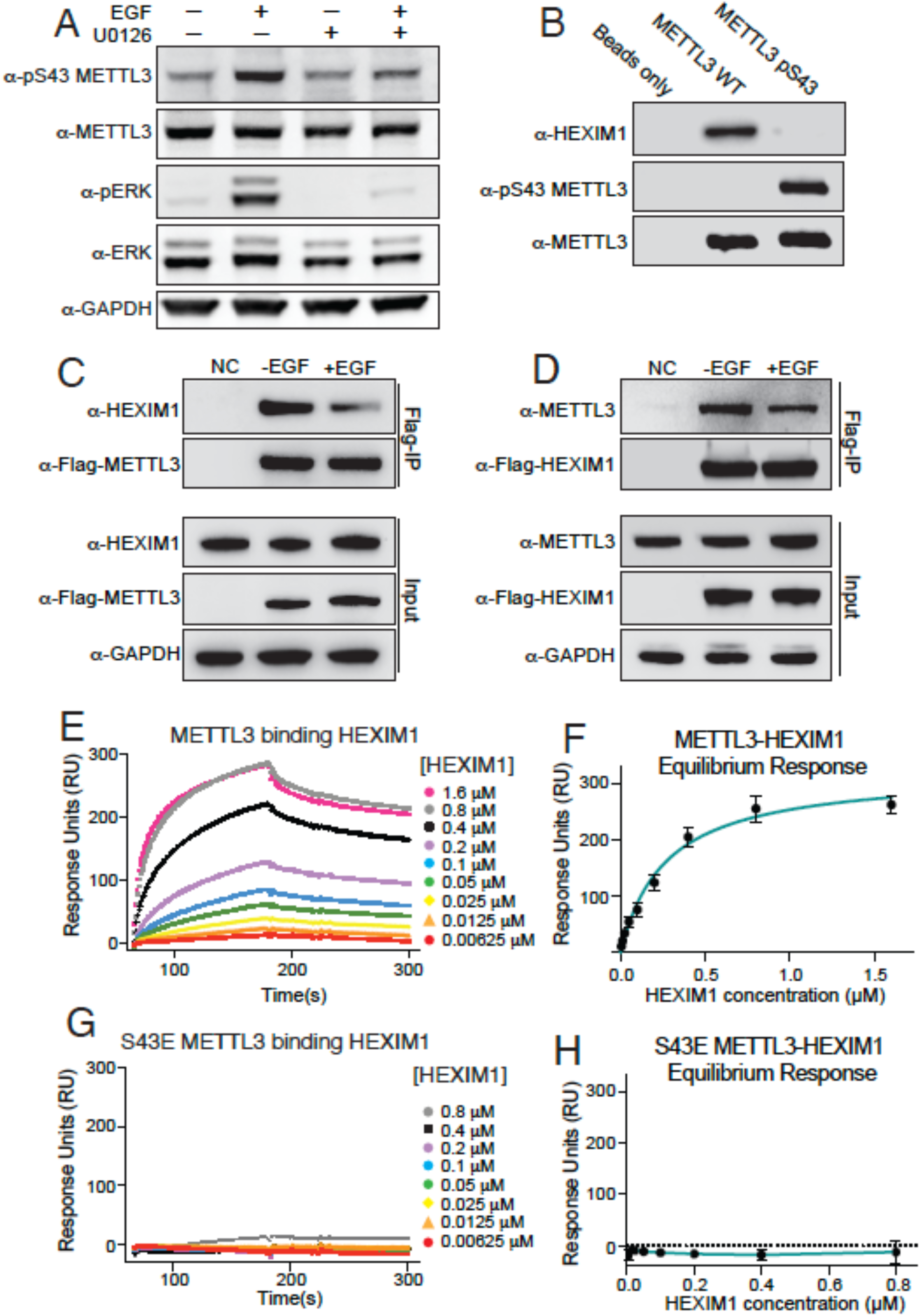
METTL3 phosphorylation prevents its interaction with HEXIM1. **(A)** Effect of U0126 on pS43-METTL3 phosphorylation induced by EGF in HeLa cells. Western blot analysis of total cell extracts using the indicated antibodies. Blots in A-D are representatives of three biological repeats. **(B)** Pull-down of endogenous HEXIM1 from HeLa cell lysates using beads containing GST-tagged unphosphorylated or pS43 METTL3, produced in bacteria. Beads alone were used as a negative control. Western blot analysis using the indicated antibodies. Blots are a representation of three biological repeats. **(C)** Immunoprecipitation of stably expressed Flag-METTL3. The top two panels represent the IP and the bottom three the input. Western blot analysis using the indicated antibodies. Wild-type HeLa cells were used as a negative control (NC) for the IP. **(D)** Immunoprecipitation of stably expressed Flag-HEXIM1. **(E, G)** Sensorgram of Surface Plasmon Resonance (SPR) kinetic analysis of HEXIM1 binding to immobilized METTL3 (**E**) or phosphomimetic S43E METTL3 (**G**). **(F, H)** Non-linear regression of D and F respectively. The graphs show mean ± SD from three biological replicates.

We hypothesized that METTL3 phosphorylation on S43 could impact the interaction between METTL3 and some of its protein partners. To test this, we generated recombinant GST-METTL3 phosphorylated at position S43 using the orthogonal translation system SepOTS (*37*). This system is based on an *Escherichia coli* strain engineered to fully incorporate phosphoserine (Sep) genetically into recombinant proteins (*37*). We validated the presence of pS43 by western blot techniques (Fig. 3B). We then used glutathione beads loaded with GST-pS43-METTL3 and the non-phosphorylated GST-METTL3 counterpart protein fusions as ‘baits’ to pull down interacting proteins from HeLa cell lysates obtained by cryomilling. We found that 7SK-interacting protein HEXIM1 specifically interacts with non-phosphorylated METTL3 (Fig. 3B). These results strongly suggest that S43 phosphorylation disrupts the interaction between METTL3 and HEXIM1.

Since EGF stimulates the phosphorylation of METTL3 on S43, we next tested the effect of EGF stimulation on METTL3/HEXIM1 interactions in a cellular context. Using cryomilled lysates from serum-starved cells or cells stimulated with EGF, we immunoprecipitated Flag-METTL3 and performed western blots for endogenous HEXIM1 (Fig. 3C). Consistent with the GST fusion pull-down experiments, we found that EGF stimulation reduced the interaction between METTL3 and HEXIM1. These findings were confirmed with reciprocal co-immunoprecipitation using Flag-HEXIM1 (Fig. 3D).

To determine if the interaction between METTL3 and HEXIM1 is direct, we employed Surface Plasmon Resonance (SPR). Purified METTL3 and HEXIM1 produced in insect cells, showed high-affinity binding with a dissociation constant (*K*_d_) of approximately 0.27 ± 0.05 μM (Fig. 3E-F and S3D-E), whereas the phosphomimetic mutant S43E abrogated the interaction (Fig. 3G-H). These results demonstrate a direct interaction between METTL3 and HEXIM1 and a negative impact of the phosphomimetic S43E residue on this interaction. Moreover, these results are consistent with the disruptive effect of pS43 on the formation of the METTL3-HEXIM1 complex.

### METTL3 is required for EGF-induced 7SK-interactome switch and transcriptional activity

To determine the impact of EGF on 7SK methylation, we pulled down m^6^A-methylated nuclear RNA from unstimulated and EGF-treated cells using an m^6^A antibody, followed by a qRT-PCR for 7SK. As shown in Fig. 4A, EGF stimulation increased the levels of m^6^A-methylated 7SK several-fold. Importantly, this effect was prevented by METTL3 depletion, indicating that METTL3 is required for EGF-stimulated methylation of 7SK (Fig. 4A). Since EGF stimulation induces 7SK methylation, and METTL3 depletion increases HEXIM1 binding to 7SK, we predicted that EGF stimulation should diminish the interaction between HEXIM1 and 7SK. As expected, EGF stimulation significantly reduced the interaction between HEXIM1 and 7SK, as shown by Northern blotting for 7SK after immunoprecipitation of HEXIM1 (left lanes of Fig. 4B). Importantly, this effect was lost in cells depleted of METTL3, indicating that the ability of EGF to induce the release of HEXIM1 from 7SK depends on METTL3 (Fig. 4B). In contrast and as expected, EGF stimulation enhanced HNRNPA2B1 binding to 7SK, an outcome that was eliminated by depletion of METTL3 (Fig. 4C). Consistent with the results shown in Fig. 2E-F, basal levels of 7SK bound to HEXIM1 increased upon METTL3 depletion, with the opposite observed for HNRNPA2B1 (Fig. 2E-F and 4B-C).

**Fig. 4.**
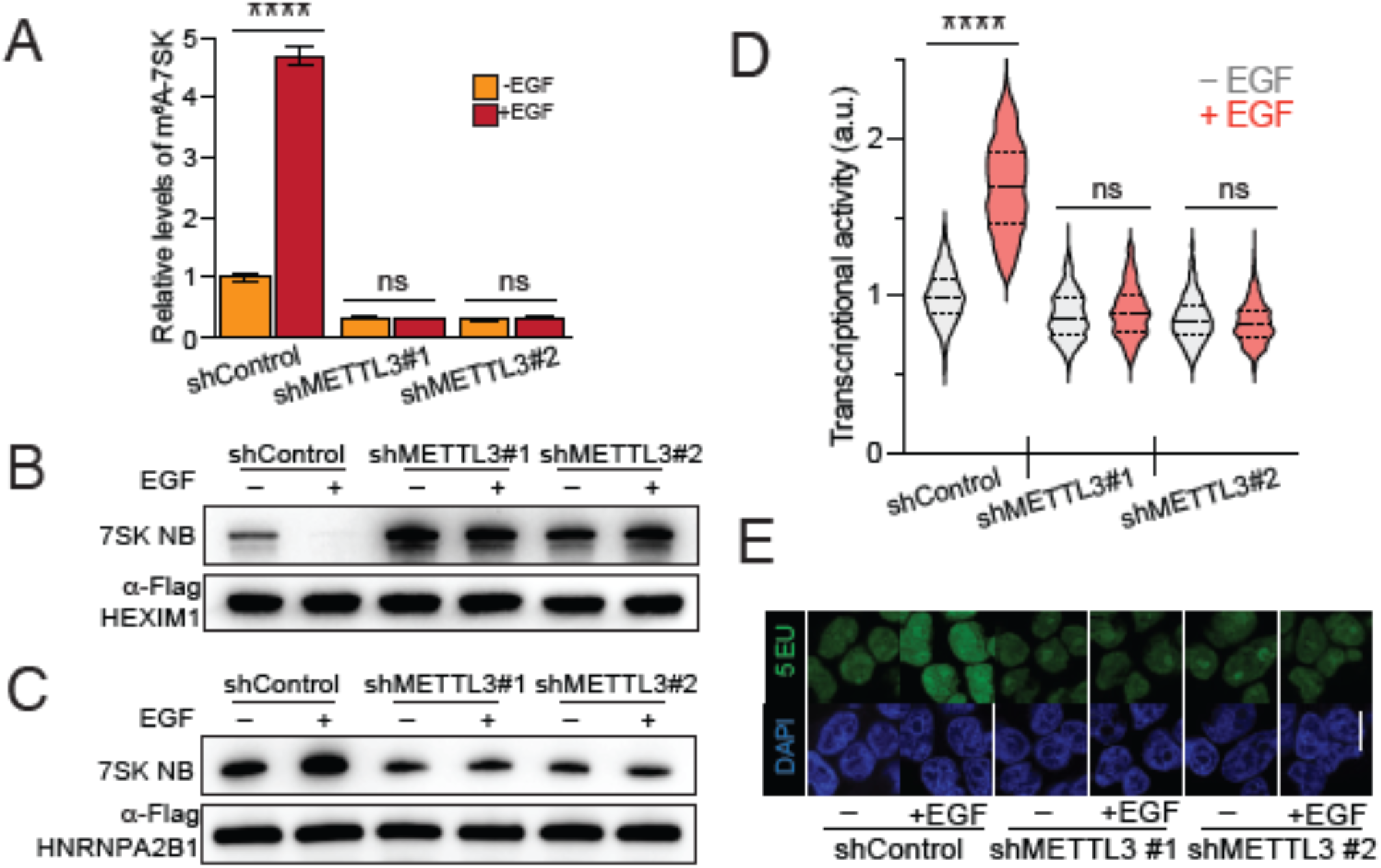
METTL3 depletion favors HEXIM1 binding to 7SK and decreases EGF-induced transcriptional activation. **(A)** m^6^A methylation levels on 7SK upon EGF stimulation in wild-type cells and cells depleted of METTL3. Nuclear m^6^A-IP followed by qRT-PCR quantification of 7SK upon EGF stimulation. A control shRNA and two independent shRNAs targeting METTL3 were stably expressed in HeLa cells. The graph shows mean ± SD from a representative experiment out of three biological replicates, with three technical replicates each. Two-way ANOVA, Tukey’s post-test, \*\*\*\**P*<1×10^−4^. **(B, C)** 7SK-protein interaction with HEXIM1 (**B**) and HNRNPA2B1 (**C**), upon EGF stimulation in control cells and cells depleted of METTL3 using two independent shRNAs. The top panels show the pull-down endogenous 7SK measured by NB and the loading immunoprecipitated corresponding proteins detected by WB. Blots are a representation of two biological replicates. **(D)** Cellular transcriptional activity upon EGF stimulation in control cells and cells depleted of METTL3 using two independent shRNAs, as measured by RNA Click-iT. Violin plots of fluorescent measurement of 600 cells per condition. Bottom: micrographs of representative cell nuclei (**E**). A representative experiment of two biological replicates, with three technical replicates each. Two-way ANOVA, Tukey’s post-test, \*\*\*\**P*<1×10^−4^.

Since it is stablished that the release of HEXIM1 from 7SK results in transcriptional elongation via the activity of free P-TEFb, we tested the functional role of m^6^A on EGF-stimulated transcription. To do this, we used the quantitative imaging technique called Click-iT, which enables the detection of global RNA transcription in cells (61). As shown in Fig. 4D-E, EGF stimulated transcription in control cells. However, the depletion of METTL3 eliminated the ability of EGF to stimulate transcription, indicating that m^6^A is required for this EGF-induced response (Fig. 4, D-E). These results indicate that m^6^A is required for the EGF-induced exchange in 7SK binding of HEXIM1 for HNRNPs, and subsequent transcriptional activity (Fig. 4B-E).

### 7SK-interactome switch depends on METTL3 phosphorylation

We decided to test whether the effect of EGF on the methylation levels of 7SK was due to the ERK activity. We found that the MEK inhibitor U0126 blocked the EGF-induced increase in 7SK methylation, indicating that ERK activity is required for this process (Fig. 5A). To determine if METTL3 phosphorylation downstream of ERK is required for 7SK methylation, we performed a loss-of-function experiment using CRISPR/Cas9 to eliminate the S43 phosphorylation site of endogenous METTL3. We obtained several positive clones carrying the S43A homozygous mutation (METTL3^S43A^). These mutated cell lines, as expected, were defective in EGF-induced S43 phosphorylation (Fig. 5B). We then used these clones to assess the impact of pS43 on the ability of METTL3 to methylate 7SK. As shown in fig. 5C, wild-type cells efficiently increased the 7SK methylation upon EGF stimulation. However, two independent METTL3^S43A^ clones had lost the EGF-induced m^6^A-methylation capacity of 7SK (Fig. 5C). These results indicate that the ERK-mediated phosphorylation of METTL3 S43 is required for the ability of the methyltransferase to methylate 7SK upon EGF stimulation.

**Fig. 5.**
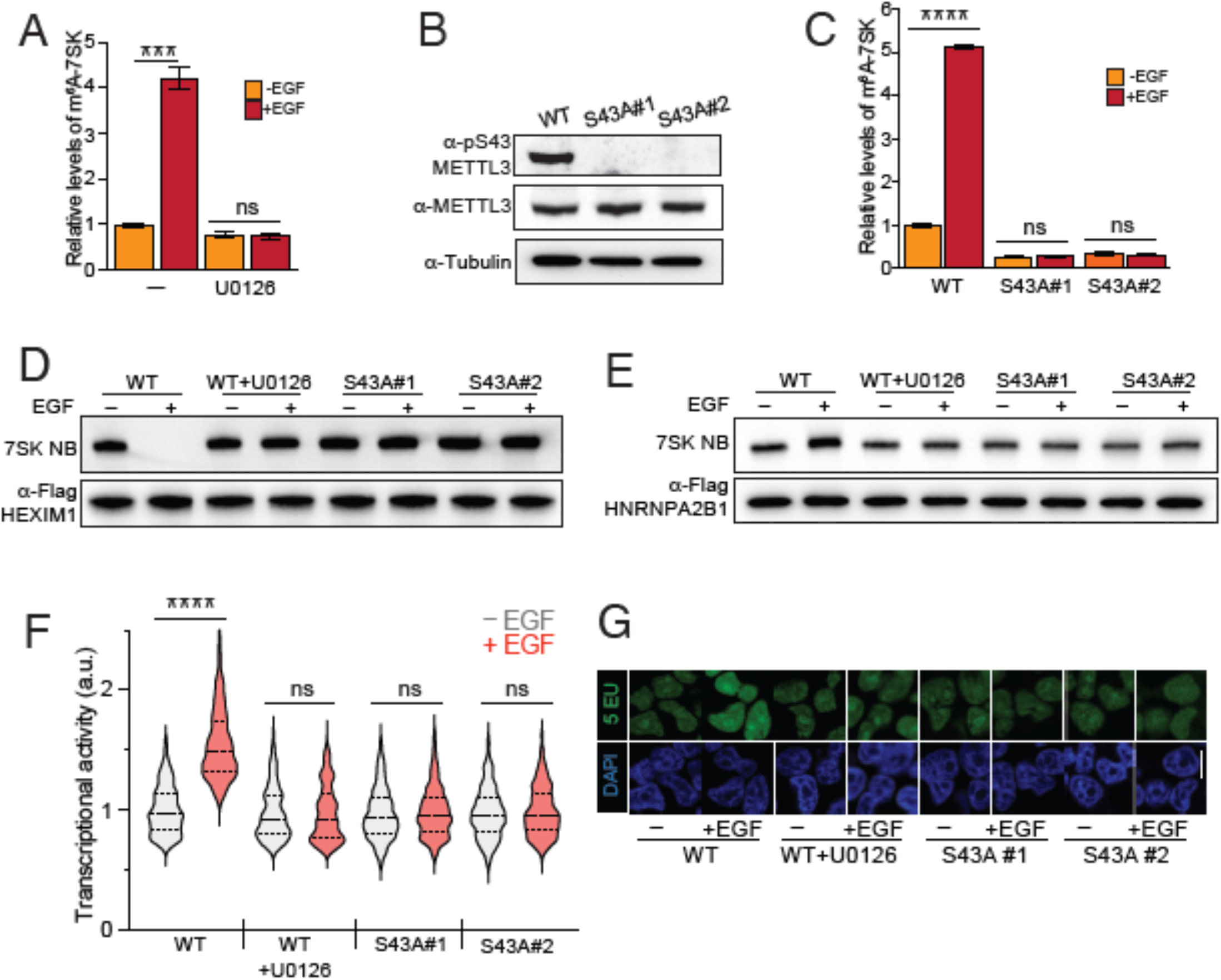
EGF transcriptional activation depends on METTL3 phosphorylation. **(A)** Effect of U0126 on the EGF-induced m^6^A methylation of 7SK. Experimental design is similar to 4A. The graph shows mean ± SD from a representative experiment out of two biological replicates, with three technical replicates each. One-way ANOVA, Tukey’s post-test, \*\*\**P*<1×10^−3^. **(B)** CRISPR/Cas9 mutagenesis of S43A of METTL3. Western blot of METTL3 pS43 upon EGF stimulation in wild-type (WT) cells and two METTL3^S43A^ homozygous clones. **(C)** Similar to A, but in this case, wild-type cells and two METTL3^S43A^ homozygous clones were used. The graph shows mean ± SD from a representative experiment out of two biological replicates, with three technical replicates each. Two-way ANOVA, Tukey’s post-test, \*\*\*\**P*<1×10^−4^. **(D, E)** 7SK-protein interaction with HEXIM1 (**D**) and HNRNPA2B1 (**E**), upon EGF stimulation in wild-type cells and two independent METTL3^S43A^ homozygous clones. Blots are representative of two biological replicates. **(F**,**G)** Effect of U0126 and mutagenesis of S43A-METTL3 on transcriptional activity induced by EGF stimulation. A representative experiment of two biological replicates, with three technical replicates each. Two-way ANOVA, Tukey’s post-test, \*\*\*\**P*<1×10^−4^.

To assess the role of METTL3 phosphorylation on the ability of 7SK of dissociate from HEXIM1 upon EGF stimulation, we immunoprecipitated HEXIM1 and assessed the levels of associated 7SK by Northern blot. As shown in Fig. 5D, U0126 blocked the negative impact of EGF on the HEXIM1/7SK interaction (Fig. 5D). Importantly, this effect was mimicked by the METTL3^S43A^ clones, which did not exhibit modulation of the HEXIM1 binding to 7SK (Fig. 5D). As expected, the impact of MEK inhibition and S43A mutagenesis on HNRNPA2B1 was the opposite of that observed with HEXIM1. Consistent with the data shown so far, inhibiting ERK activation prevented the enhancement of HNRNPA2B1 binding to 7SK upon EGF stimulation, an effect that was phenocopied by eliminating the S43 phosphorylation site in the METTL3^S43A^ clones (Fig. 5E).

We further tested if the EGF requirement of m^6^A for transcriptional activity relies on the ability of EGF to induce ERK-mediated phosphorylation of METTL3. As anticipated, the effect of EGF on transcriptional activity was blocked by both the MEK inhibitor U0126 and by the elimination of the S43 METTL3 phosphorylation site (Fig. 5F-G). Together, these results demonstrate that METTL3 and its ERK-mediated phosphorylation of S43 are required for the EGF-induced 7SK-m^6^A methylation and the subsequent transition from 7SK-bound HEXIM1 to HNRNPs complexes, resulting in enhanced transcriptional activity.

### 7SK-methylation is required for interactome switch and transcriptional activity

To test if the methylation of 7SK is directly responsible for the interactome switch and transcriptional activity, we decided to mutate the adenosines corresponding to the predicted methylation sites described above. We used CRISPRi to inactivate the expression of endogenous 7SK (*38*). We then used a lentiviral expression system to express exogenous 7SK, wild type or m^6^A-mutant, under the RNA Pol III U6 promoter (Fig. 6A). As shown in Fig. 6B, elimination of the predicted site 1 did not affect the levels of m^6^A on 7SK compared to the expression of exogenous wild type version of 7SK, while elimination of the sites 2-5, corresponding to the HNRNPs binding region, almost completely eliminated the levels of m^6^A on 7SK (Fig. 6B). Based on the experiments presented this far, we predicted that elimination of the m^6^A sites would result in the inability of 7SK to release HEXIM1 and P-TEFb and therefore unable to stimulate transcription upon EGF stimulation. As expected, wild type 7SK efficiently released HEXIM1 upon EGF stimulation, while the m^6^A-mutant version of 7SK was inert to the stimulation by the growth factor (Fig. 6C). Moreover, 7SK depletion by CRISPRi showed an enhanced basal transcriptional activity compared to control wildtype cells, consistent with its role in sequestering the transcription elongation factors (Fig. 6D-E). However, the transcriptional activity of 7SK depleted cells failed to be regulated by EGF, demonstrating that the EGF effect on transcription depend on 7SK (fig. 6D-E). Importantly, reintroduction of 7SK wildtype rescued the level of basal transcriptional levels and the regulation by EGF stimulation, while m^6^A-mutant 7SK showed a decreased and unresponsive transcriptional activity (Fig. 6D-E). These results demonstrate that the methylation of 7SK is required to release HEXIM1/P-TEFb and promote transcription upon EGF stimulation.

**Fig. 6.**
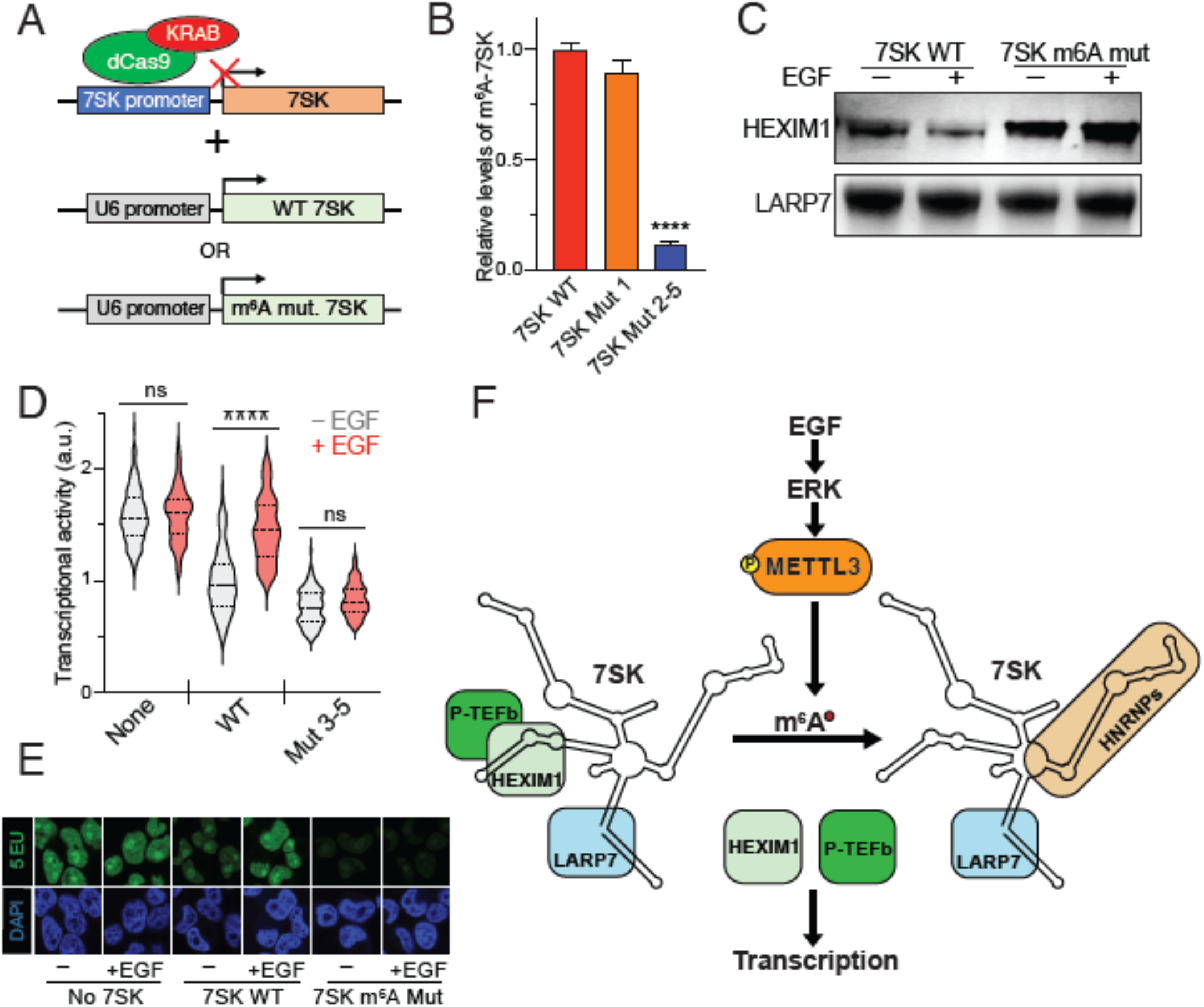
7SK m6A methylation is required for HEXIM1 dissociation and transcriptional activation downstream EGF signaling. **(A)** CRISPRi was used to downregulate the expression of 7SK. Once cells were depleted of endogenous 7SK, lentiviral vectors were used to stably express exogenous wildtype or m^6^A mutant 7SK under the U6 promoter. **(B)** Immunoprecipitation of m^6^A methylated RNA followed by qRT-PCR of 7SK of cells depleted of 7SK and transduced with wildtype 7SK (red bar), 7SK mutant for site #1 of m^6^A (orange bar), and 7SK mutant for sites #2-5 (blue bar). **(C)** Flag-LARP7 immunoprecipitation of CRISPRi 7SK plus wildtype 7SK or 7Sk mutant for sites #2-5 upon EGF stimulation. Western blot for HEXIM1 and LARP7 using the indicated antibodies. **(D, E)** Effect of mutagenesis of m^6^A sites of 7SK on transcriptional activity induced by EGF stimulation. A representative experiment of two biological replicates, with three technical replicates each. Two-way ANOVA, Tukey’s post-test, ****P<1×10-4. **(F)** Model. The work presented here is summarized in this model. Upon EGF stimulation the downstream effector ERK kinase is activated and phosphorylates METTL3 at position S43. This phosphorylation results in the release of METTL3 from its association with HEXIM1, thus allowing the methylation of the snRNA 7SK. Subsequently, m^6^A methylation of 7SK results in its association with HNRNP proteins and release of the HEXIM1/P-TEFb complex. Free P-TEFb is then available to induce transcription activation.

## Discussion

In this report we uncovered 7SK as a novel substrate of METTL3/m^6^A methylation and a new mechanism of transcriptional control downstream signaling pathways (fig. 6F). Based on our results, we propose a model whereby, in unstimulated conditions, METTL3 is sequestered by HEXIM1. Once cells get growth promoting signals, such EGF, ERK-mediated phosphorylation of METTL3 results in its dissociation from HEXIM1 and consequent methylation of 7SK. Once 7SK is methylated, it releases HEXIM1 and the P-TEFb complex and in turn associates with HNRNP proteins. Finally, the release of P-TEFb results in enhanced transcriptional activity (Fig. 6F).

7SK is an abundant non-coding RNA evolutionarily conserved in vertebrates, and homologs have been found in insects and other animals (*39*). The secondary and tertiary structures of 7SK are not entirely elucidated; however, 7SK is known to fold into several hairpins recognized by distinct binding proteins. 7SK associates with a subset of HNRNPs when it is not associated with HEXIM1. Several post-translational modifications on the 7SK-associated proteins have been proposed to increase or decrease their affinity for 7SK. However, the mechanism for exchanging between the two protein complexes is not well understood. Our identification of m^6^A methylation of 7SK provides a novel regulatory mechanism for the 7SK transition from an anti to a pro-proliferative state. In this novel mechanism, the RNA itself is modified upon cellular stimulation by growth factors (Fig. 6F). This novel growth factor-induced methylation by METTL3 of the mature 7SK, an RNA Pol III transcript, contrasts with the known co-transcriptional activity of METTL3 on RNA Pol II transcripts such as mRNA and primary miRNAs (*24*), thus increasing the type of RNA target molecules of METTL3.

Our results show that m^6^A is localized in the hairpin recognized by HNRNPs. A recent iCLIP experiment of HNRNPR shows a significant overlap between its binding regions and the m^6^A motifs (*36*). Importantly, the “HNRNP hairpin” is opposite to the one recognized by HEXIM1. Thus, when the HEXIM1/P-TEFb complex is bound to 7SK, it is likely that the HNRNP hairpin is free and available for METTL3 recognition and methylation. It has been shown that HNRNPC, HNRNPG, and HNRNPA2B1 bind methylated RNA indirectly, through a structural switching mechanism (*40-42*). Thus, it is plausible that all the 7SK-interacting HNRNP proteins, none of which have specialized m^6^A binding domains seen in YTHD readers, use a structural switching mechanism to bind methylated 7SK. It would be interesting to know if the HNRNPs that recognize methylated 7SK represent redundancy, respond to different stimuli, or are regulated in a cell type-specific manner.

METTL3 phosphorylation has been associated with changes in its subcellular localization and probably activity (*43*), demonstrating this as one regulatory mechanism of m^6^A methylation. Here we show that a major signaling pathway induce functional METTL3 phosphorylation to regulate transcriptional activity. We found that ERK phosphorylates METTL3 at S43 upon EGF stimulation. This phosphorylation releases METTL3 from HEXIM1 binding allowing methylation of 7SK. Others have recently shown the importance of S43 phosphorylation in different contexts (*43, 44*), supporting the relevance of this phosphorylation site for METTL3 activity.

As the EGF signaling pathway induced the activation of METTL3, resulting in transcription activity through a 7SK-mediated mechanism, it is important to note that, once activated, METTL3 could have additional downstream consequences for the newly transcribed mRNAs either to promote translation, splicing, or degradation. Importantly, as EGF and other signaling pathways are altered in diseases such as cancer, it would be important to approach m^6^A as a downstream effector of such dysregulations. Finally, since 7SK has been shown to participate in cell differentiation, it would be interesting to assess the mechanism overall presented here in the context of differentiation of progenitors or stem cells.

## Acknowledgments

We thank M. Lemmon and S. F. Tavazoie for reading earlier versions of this manuscript. We acknowledge support from our colleagues at the Yale Cancer Biology Institute. We thank the Yale West Campus Imaging Core for the support and assistance in this work. We thank C. Osuji and P. Rahmani for their technical support.

## Funding

National Institutes of Health grant R35GM138185 (CRA, AWD, HL)

National Institutes of Health grant R01GM137031 (YL)

National Institutes of Health grant R01CA248532(DEK)

National Institutes of Health grant R01CA244634 (HG)

CRA is a Louis Goodman and Alfred Gilman Yale Scholar.

## Author contributions

(based on https://credit.niso.org/) Conceptualization: CRA

Formal Analysis: MPP Methodology: MPP, AWD

Investigation: MPP, AWD, HL, WL, BH, QL, HG Visualization: MPP, CRA

Supervision: DEK, YL, CRA Writing – original draft: MPP, CRA

## Competing interests

The authors declare no competing interests.

## Data and materials availability

Constructs will be deposited in Addgene or available upon request with the corresponding MTA.

## Notes

### Competing Interest Statement

The authors have declared no competing interest.

### Summary of Updates

The title of the manuscript was changed from "7SK methylation Promotes Transcriptional Activity Upon EGF Stimulation" to "7SK methylation Promotes Transcriptional Activity"

